# Phasic inhibition of dopamine neurons is an instrumental punisher

**DOI:** 10.1101/2020.11.25.399220

**Authors:** Constance Yunzhi Peng, Philip Jean-Richard-dit-Bressel, Sophia Gilchrist, John M. Power, Gavan P. McNally

**Author notes:** Correspondence to: Gavan P. McNally, School of Psychology, UNSW, Sydney, 2052, Australia, e., p.

## Abstract

It is well established that the activity of VTA dopamine neurons is sufficient to serve as a Pavlovian reinforcer but whether this activity can also serve as instrumental reinforcer is less well understood. Here we studied the effects of optogenetic inhibition of VTA dopamine neurons in instrumental conditioning preparations. We show that optogenetic inhibition of VTA dopamine neurons causes a response-specific, contingency-sensitive suppression of instrumental responding. This suppression was due to instrumental response, not Pavlovian stimulus, learning and could not be attributed to deepened instrumental extinction learning. These effects of optogenetic inhibition of VTA dopamine neurons on instrumental responding are formally similar to the effects of aversive events in instrumental preparations and show that optogenetic inhibition of VTA dopamine neurons is sufficient to serve as an instrumental punisher.

---

The activity of VTA dopamine neurons and consequent release of dopamine in the striatum serves multiple roles in associative learning. In Pavlovian learning, VTA dopamine neurons signal reward prediction errors instructing Pavlovian association formation (Schultz, 1997; Schultz, 1999, 2002, 2007; Schultz, 2017; Schultz & Dickinson, 2000). Increases in the activity of VTA dopamine neurons signal positive prediction error and instruct increments in associative strength whereas decreases in their activity signal negative prediction errors and instruct decreases in associative strength. A variety of lines of evidence, notably single unit recordings and optogenetic manipulation of these neurons, supports this role (Chang et al., 2016; Chang et al., 2018; Langdon et al., 2017; Saunders et al., 2018; Sharpe et al., 2017; Steinberg et al., 2013). For example, brief experimenter-administered VTA dopamine neuron excitation supports Pavlovian appetitive contextual (Tsai et al., 2009) or discrete conditioned stimulus (CS) appetitive learning (Saunders et al., 2018) whereas inhibition is sufficient to imbue a CS with inhibitory properties (Chang et al., 2018) and a context with aversive motivational properties (Danjo et al., 2014; Ilango et al., 2014; Tan et al., 2012).

The role of VTA dopamine neurons in instrumental learning is more complex and less clear. A common assumption is that instrumental learning is also instructed by a bidirectional reinforcement signal that supports reward (when positive) or punishment (when negative) (Sutton & Barto, 2018). These are modern versions of Thorndike’s Law of Effect (Thorndike, 1898). VTA dopamine neurons could serve this role (Sutton & Barto, 2018) but so too could other neurotransmitters such as serotonin (Daw et al., 2002). Consistent with a role for excitation of VTA dopamine neurons in instrumental reinforcement, studies of intracranial optical self-stimulation whereby excitation of VTA dopamine neurons is contingent on a nominally instrumental behavior such as lever pressing or nosepoking identify potent reinforcing effects of this stimulation (Ilango et al., 2014; Kim et al., 2012; Steinberg et al., 2014; Witten et al., 2011) (but see (Adamantidis et al., 2011). However, the instrumentality of behavior in these studies is often unclear, especially given the well documented roles for VTA dopamine neurons in Pavlovian appetitive learning (see above) and Pavlovian sign tracking behavior towards levers (Flagel et al., 2011; Saunders et al., 2018) that could equally well explain approach and lever pressing behaviors in many optical self-stimulation studies.

Even less clear is whether inhibition of VTA dopamine neurons is sufficient to serve as instrumental punisher. Punishment involves the learning of Response–Outcome associations that selectively suppress punished, but not other behaviors, as a function of the contingency between the instrumental response and the outcome (Bolles et al., 1980; Bolles et al., 1975; Goodall, 1984; Jean-Richard-Dit-Bressel et al., 2018). Punishment is neither equivalent nor reducible to Pavlovian fear learning (Bolles et al., 1980; Goodall, 1984; Goodall & Mackintosh, 1987; Jean-Richard Dit Bressel & McNally, 2014, 2015; Jean-Richard-Dit-Bressel et al., 2019; Jean-Richard-Dit-Bressel & McNally, 2016; Killcross et al., 1997). Certainly, rodents will readily learn to avoid a location that is associated with direct (Danjo et al., 2014; Ilango et al., 2014) or indirect (Jhou et al., 2013; Stamatakis & Stuber, 2012; Tan et al., 2012) inhibition of VTA dopamine neurons. But, whether avoidance in these studies is driven by Pavlovian or instrumental learning and whether such avoidance was due specifically to instrumental punishment learning is unknown. Further complicating interpretation of the effects of VTA dopamine inhibition in instrumental learning is that dopamine neurons have roles not just in reinforcement but also in the initiation of instrumental behaviors (Fischbach-Weiss et al., 2018; Hamid et al., 2015; Mohebi et al., 2019; Nicola, 2010; Wassum et al., 2012). Response-contingent inhibition of VTA dopamine neurons could suppress instrumental behavior not because it is an effective instrumental punisher but instead because it prevents initiation or maintenance of the instrumental response or even because it reduces the reinforcing potency of the instrumental reward (see (Fischbach-Weiss et al., 2018; Jean-Richard-Dit-Bressel et al., 2018) for discussion).

Here we used optogenetic approaches in TH::Cre transgenic rats to ask whether VTA dopamine neuron inhibition is sufficient to serve as instrumental punisher. The questions of interest were whether photoinhibition contingent on instrumental lever-pressing suppressed lever-pressing in a response-specific manner and whether suppression was sensitive to the contingency and duration of photoinhibition. We also sought to address the role of instrumental response learning versus Pavlovian stimulus learning in suppression, and how such suppression relates to instrumental extinction learning. Our results show that optogenetic inhibition of VTA dopamine neurons is sufficient to serve as an instrumental punisher and that the effects of this inhibition are formally similar to the effects of other instrumental punishers.

## Method

### Subjects

Subjects were experimentally naive male SD-TH-Cre^tm1sage^ +/− (TH::Cre+) rats or their SD-TH-Cre^tm1sage^ −/− (TH::Cre−) littermates (Horizon Discovery) (300-500g) obtained from Animal Resources Centre. They were housed in groups of four in ventilated plastic cages and maintained on a 12-hour light–dark cycle (lights on 7:00a.m.). Food (Gordon’s Rat and Mouse Food) and water were freely available until three days before the start of all behavioral training. All procedures were approved by the Animal Care and Ethics Committee at the University of New South Wales and conducted in accordance with the National Institutes of Health (NIH) Guide for the Care and Use of Laboratory Animals (NIH Publications No. 80–23, revised 1996).

### Apparatus

All behavioral procedures were conducted in a set of eight identical experimental chambers [24 cm (length) × 30 cm (width) × 21 cm (height); Med Associates Inc.]. Each chamber was enclosed in sound- and light-attenuating cabinets [59.5 cm (length) × 59 cm (width) × 48 cm (height)] and fitted with fans for ventilation and background noise. The chambers were constructed of a Perspex rear-wall, ceiling and hinged front-wall, and stainless-steel sidewalls. The chamber floors consisted of stainless-steel rods (4 mm in diameter) spaced 15 mm apart. Each chamber stood 35 mm above a tray of corncob bedding. An external automatic hopper dispensed a 45 g grain pellets (Able Scientific) into a recessed magazine (3 cm in diameter) within a 5 × 5 cm hollow in the right-side chamber wall. Infrared photocells detected entries into the magazine. There were two retractable levers on the right-side chamber wall, on either side of the magazine. An LED driver with an integrated rotary joint (625nm wavelength; Doric Instruments) was suspended above the chamber. All chambers were connected to a computer with Med-PC IV software (Med Associates Inc.), which controlled all required manipulandum and recorded all required variables.

### Surgery

Rats were anaesthetized via an intraperitoneal injection of a mixture of 1.3 mL/kg ketamine (100 mg/mL; Ketapex; Apex Laboratories) and 0.2 mL/kg muscle relaxant, xylazine (20 mg/mL; Rompun; Bayer). They were shaved to expose the skin surface of the head and placed in the stereotaxic apparatus (Model 942, David Kopf Instruments) with the incisor bar maintained at ~3.3 mm below horizontal to achieve a flat skull position. Subjects then received a subcutaneous injection of 0.05mL carprofen (50mg/mL, Rimadyl, Pfizer) and bupivacaine (0.5%, Marcaine, Cenvet) under the surface of the incision site. A hand-drill was used to make two craniotomies above the VTA. Following incision, metal screws were positioned around the craniotomies and attached to the skull. A 23-gauge, cone tipped 5μL stainless steel injector (SGE Analytical Science) connected to an infusion pump (UMP3 with SYS4 Micro-controller; World Precision Instruments) was used to infuse 0.75μl/hemisphere of AAV vector into the VTA over 3 minutes. The viral vector, AAV5-EF1α-DIO-eNpHR3.0-eYFP (4 × 10^12^ vp/mL) or AAV5-EF1α-DIO-eNpHR3.0-mCherry (4 × 10^12^ vp/mL), was obtained from UNC Vector Core (University of North Carolina). The coordinates used were AP: −5.40, ML: ± 2.25, DV: 8.2 in mm from bregma, at a 10 degree angle (Paxinos & Watson, 2007). The needle was left in place for 7 minutes to allow for diffusion of the vector and reduce spread up the infusion tract. Hand fabricated cannulae were implanted bilaterally above the VTA according to the coordinates: AP −5.40, ML ± 2.25, D-V −7.2 in millimetres, at a 10 degree angle (Paxinos & Watson, 2007). The cannuale were secured using dental cement (Vertex Dental) anchored to the screws and skull. After surgery, rats received intramuscular injection of 0.2 mL of procaine penicillin (150 mg/mL; Benicillin; Troy Laboratories) and 0.2 mL of cephazolin sodium (100 mg/mL; AFT Pharmaceuticals). Daily monitoring of weight and behavioral changes for all rats was performed until the end of the behavioral procedures. Rats recovered for a minimum of two weeks prior to the start of the experimental procedure and were monitored daily.

### Procedure

#### Experiment 1: Photoinhibition of VTA dopamine neurons

Brain slices were prepared from TH-Cre+ rats 5 - 8 weeks after they received AAV-EF1α-DIO-eNpHR3.0-EYFP to the VTA. Rats were deeply anesthetised with 5% isoflourane gas and decapitated. Brains were rapidly extracted and placed in ice-cold cutting artificial cerebral spinal fluid (ACSF) composed of 95 mM NaCl, 2.5 mM KCl, 30 mM NaHCO_3_, 1.2 mM NaH_2_PO_4_, 20 mM HEPES, 25 mM glucose, 5 mM ascorbate, 2 mM thiourea, 3 mM sodium pyruvate, 0.5 mM CaCl_2_, and 10 mM MgSO_4_ for 3-4 minutes. Coronal slices (300 μm) that included the VTA were prepared using a vibratome (Model VT1200, Leica, Wetzlar, Germany) and incubated for 10 min in a 30°C recovery ACSF, containing equimolar N-Methyl-D-glucamine in place of NaCl. Slices were then transferred to a Braincubator (#BR26021976, Payo Scientific, Sydney, Australia) where they were maintained at 16°C in ACSF containing 95 mM NaCl, 2.5 mM KCl, 30 mM NaHCO_3_, 1.2 mM NaH_2_PO_4_, 20 mM HEPES, 30 mM glucose, 5 mM ascorbate, 2 mM thiourea, 3 mM sodium pyruvate, 2 mM CaCl_2_ and 2 mM MgSO_4_, until recording. All solutions were pH adjusted to 7.3-7.4 with HCl or NaOH and gassed with carbogen (95% O_2_ - 5% CO_2_).

Brain slices were transferred to a recording chamber, maintained at 30°C and continuously perfused with oxygenated standard ACSF (containing in mM; 124 NaCl, 3 KCl, 1.2 NaH_2_PO_4_, 26 NaHCO_3_, 10 glucose, 1.3 MgCl_2_ and 2.5 CaCl_2_.). Targeted whole-cell patch-clamp recording were made from soma of EYFP+ neurons using a microscope (Zeiss Axio Examiner D1) equipped with 20x water immersion objective (1.0 NA), LED fluorescence illumination system (pE-2, CoolLED) and an EMCCD camera (iXon+, Andor Technology). Patch pipette (3–5 MΩ) were fabricated from borosilicate glass (GC120TF-4; Warner Instruments) using a two-stage vertical puller (PC-10; Narishige) and filled with an internal solution containing (in mM); 130 potassium gluconate, 10 KCl, 10 HEPES, 4 Mg_2_-ATP, 0.3 NA_3_-GTP, 0.3 EGTA, and 10 phosphocreatine disodium salt (pH adjusted to 7.25 with KOH). Electrophysiological recordings were amplified using a Multiclamp 700B amplifier (Molecular Devices) filtered at 6 kHz and digitised at 20 kHz with a multifunction I/O device (PCI-6221, National Instruments). Recordings were controlled and analysed offline using Axograph (Axograph, Sydney, Australia). The liquid junction potential (~ 9 mV) was not compensated for. The locations of all recorded cells were mapped according to the Rat Brain Atlas (Paxinos & Watson, 2013).

Series resistance and membrane resistance were calculated from current response to a voltage step from –65 to −70mV using in built routines in Axograph. Membrane time constant was determined by fitting an exponential to the voltage response to a small hyperpolarising current. The AP half-width (width at 50% of the peak) was measured relative to the AP threshold. eNpHR3.0 was stimulated using orange light (GYR LED bandpass filtered 605/50 nm) delivered through the objective. To determine the amplitude of light-evoked hyperpolarisation neurons were maintained at subthreshold membrane potentials using hyperpolarizing current injections. To determine the effect of photoinhibition on the AP firing frequency, neurons that had spontaneous firing frequencies < 2 Hz were induced to fire via depolarising current injection, or via trains of depolarising current injections. When possible, protocols were repeated 2-5 times and the results averaged. Data were excluded if the series resistance was > 25 MΩ or more than 200 pA was required to maintain the neuron at –65 mV.

#### Experiment 2: Contingency and duration sensitivity of photoinhibition

In this and remaining experiments, commencing at least two weeks after surgery and persisting for the duration of the behavioral procedures, rats received daily access to 10-15g of food and unrestricted access to water in their home cages. Rats were food deprived for two days in their home cages then received one experimental session per day. They were first placed in the experimental chambers for two 1-hour magazine training sessions, where lever-pressing on either of two concurrently-presented levers (left and right) was reinforced with grain pellets on a fixed ratio-1 (FR-1) schedule. In these sessions, a lever retracted once it was pressed 25 times. Houselights were on throughout. Any rats that did not acquire lever pressing after two days of magazine training were manually shaped until lever pressing was acquired.

In following lever-press training sessions, levers were then presented individually in an alternating pattern so that one lever was extended for 5 min while the other lever was retracted. After 5 min, the extended lever was retracted and the retracted lever was extended, such that each lever was always presented on its own. This alternation occurred throughout the 40-min session for a total of 8 trials, 4 for each lever. Lever-pressing on either lever was reinforced with pellets on a 30-sec variable interval (VI-30sec) schedule. Rats were given 9 days of lever-press training, with rats being habituated to the tethering procedure for the last 5 days of training.

Rats (TH::Cre+ n = 12, TH::Cre− n = 12) were connected to bilateral patch cables outputting at least 9mW of 625nm light, measured through an unimplanted fiber optic cannula before each session commenced. Rats continued to receive pellets on a VI-30sec schedule through all test and rest sessions. Test sessions were identical to the lever pressing training sessions, except that responses on one lever, designated as “punished”, was immediately followed by 2, 5, or 10-sec photoinhibition on an FR-1, FR-5, or FR-10 schedule (9 test sessions in total). The other lever, designated “unpunished” lever, was not paired with photoinhibition. Each test session was followed by a rest session (8 rest sessions in total), where light delivery was suspended for both levers. The designated punished lever and whichever lever was presented first (left or right) were counterbalanced between sessions and between rats.

#### Experiment 3: Response learning versus stimulus learning

TH::Cre+ rats were randomly assigned as Master (n = 8) and paired with another “Yoked” rat (n = 8). They received the same training and test session procedures as for Experiment 1 except that for the Master rats, a designated lever was “punished” with a 5-sec 625nm LED presentation on a FR-1, FR-5, and FR-10 schedule (3 test sessions in total). Each yoked rat received a LED delivery at the same time as its paired Master rat (while the same lever was extended), regardless of the yoked rat’s behavior. There was a total of 8 trials in each 40-minute session (four punished and four unpunished trials). All rats continued to receive pellets on a VI-30sec schedule throughout. The designated punished lever and which lever was presented first was both counterbalanced between sessions and between rats, but each Master and Yoked pair always had the same lever configuration.

#### Experiment 4: Effects on instrumental extinction learning

Rats (TH::Cre+ n = 8, TH::Cre− n = 8) received the same training as Experiment 1 prior to two cycles of extinction, testing, and retraining. They rats were connected to bilateral patch cables outputting at least 9mW of 625nm light, measured through an un-implanted fiber optic cannula before session commencement. During the extinction sessions, rats underwent four 15-min sessions of instrumental appetitive extinction training where the delivery of pellets was suspended and 5-sec 625nm LED stimulation was delivered on a FR- 1 schedule. Rats then underwent a 15-minute test session where both the delivery of pellets and LED light were suspended. Following extinction, rats then received four 30-minute re-acquisition sessions where the delivery of pellets was reinstated. Rats then underwent another four extinction with photoinhibition sessions, one test session and a final re-acquisition session.

### Histology

Cannulae and viral placements were verified at the end of behavioral procedures. Rats were injected intraperitoneally with sodium pentaobarbital (100mg/kg; Lethabarb) and transcardially perfused with 0.9% saline, 1% sodium nitrite solution, 360μl/L heparin and 4% buffered paraformaldehyde (PFA). Brains were postfixed and then placed in 20% sucrose solution for 24 to 48 hours. A cryostat (Leica Microsystems) was used to collect 40μm coronal sections of the VTA, which were then preserved in phosphate-buffered azide (0.1% sodium azide) at 4°C before immunohistochemistry.

We have previously shown that Cre−dependent AAV expression is limited to TH+ neurons in the VTA of the TH-Cre rat (Liu et al., 2020). Here eYFP and mCherry labelling was detected using single-colour peroxidase immunohistochemistry. Free floating sections were washed for 30 minutes in 0.1M phosphate buffer (PB; pH 7.4), followed by 50% ethanol for 30 minutes, and then 3% hydrogen peroxide diluted in 50% ethanol for 30 minutes. Sections were then incubated for 30 minutes with 5% normal horse serum (NHS) diluted in PB, then placed in 1:2000 donkey anti-GFP (Invitrogen; A10262) or donkey anti-mCherry diluted in 0.3% Triton-X, 2% NHS, and 0.1 M PB, pH 7.4, and incubated at 4°C for 48 hours. Sections were washed three times for 20 minutes each (PB, pH 7.4), incubated in 1:2000 biotinylated donkey anti-chicken (Jackson ImmunoResearch Laboratories; 703 065 155) (diluted in a solution of 0.3% Triton-X, 2% NHS, and 0.1 M PB, pH 7.4), overnight at room temperature. The sections were then washed in PB, pH 7.4, and incubated in avidin biotin (ABC reagent; Vector Elite kit 6 μl/ml avidin and 6 μl/ml biotin) diluted with PB containing 0.2% Triton, pH 7.4. eYFP-IR or mCherry-IR were identified using a DAB (D5637-56, Sigma) reaction. Immediately before this, sections were washed twice in PB, pH 7.4, and once in 0.1M acetate buffer (pH 6.0). Sections were incubated in DAB solution (0.025% DAB, 0.04% ammonium chloride, and 0.2% D-glucose in 0.1M acetate buffer, pH 6.0). The peroxidase reaction was catalysed by 0.2 μl/ml of glucose oxidase, and then were rinsed with 0.1M acetate buffer. Fiber placements and expression of eYFP-IR or mCherry for each rat were verified and photographed using a transmitted light microscope (Olympus BX51). Only animals with fiber placements over the VTA and AAV expression in the VTA were included in the analyses.

### Data Analysis

Responses on the two levers were recorded continuously throughout the sessions. These were used to compute within-subject punishment ratios in the form A/(A+B), where A and B were total presses on punished and unpunished levers, respectively. The same ratio was also used to calculate lever preference for Experiment 2, with A and B representing total presses on L1 and L2. Ratios of 0.5 indicate no difference in pressing between the two levers, ratios less than 0.5 indicate suppression of A relative to B, and ratios greater than 0.5 indicate elevation of A relative to B. These data were analysed via between × within subject ANOVA.

## Results

### Experiment 1: Photoinhibition of VTA dopamine neurons

Whole-cell patch-clamp recordings from made from eNpHR3.0-expressing VTA neurons *in vitro* from TH::Cre+ rats to confirm the inhibitory nature of the photostimulation and its effectiveness across the durations to be used in later behavioral experiments (Figure 1A,B). We have previously reported that eNpHR3.0 expression is specific to VTA TH+ neurons in this rat line (Liu et al., 2020). The recorded neurons (12 neurons from 3 rats) were mostly spontaneously active. The membrane resistance, AP-half width and membrane time constant, were 218 ± 168 MΩ, 1.1 ± 0.4 ms, and 39.7 ± 9.8 ms respectively (mean ± SD). Presentation of orange (605 nm) light produced a rapid-onset hyperpolarisation that persisted for the duration of the light (Figure 1C). The light-evoked hyperpolarisation reliably supressed neuronal firing throughout the light pulse regardless of the duration of photostimulation (Figure 1D,E).

**Figure 1.**
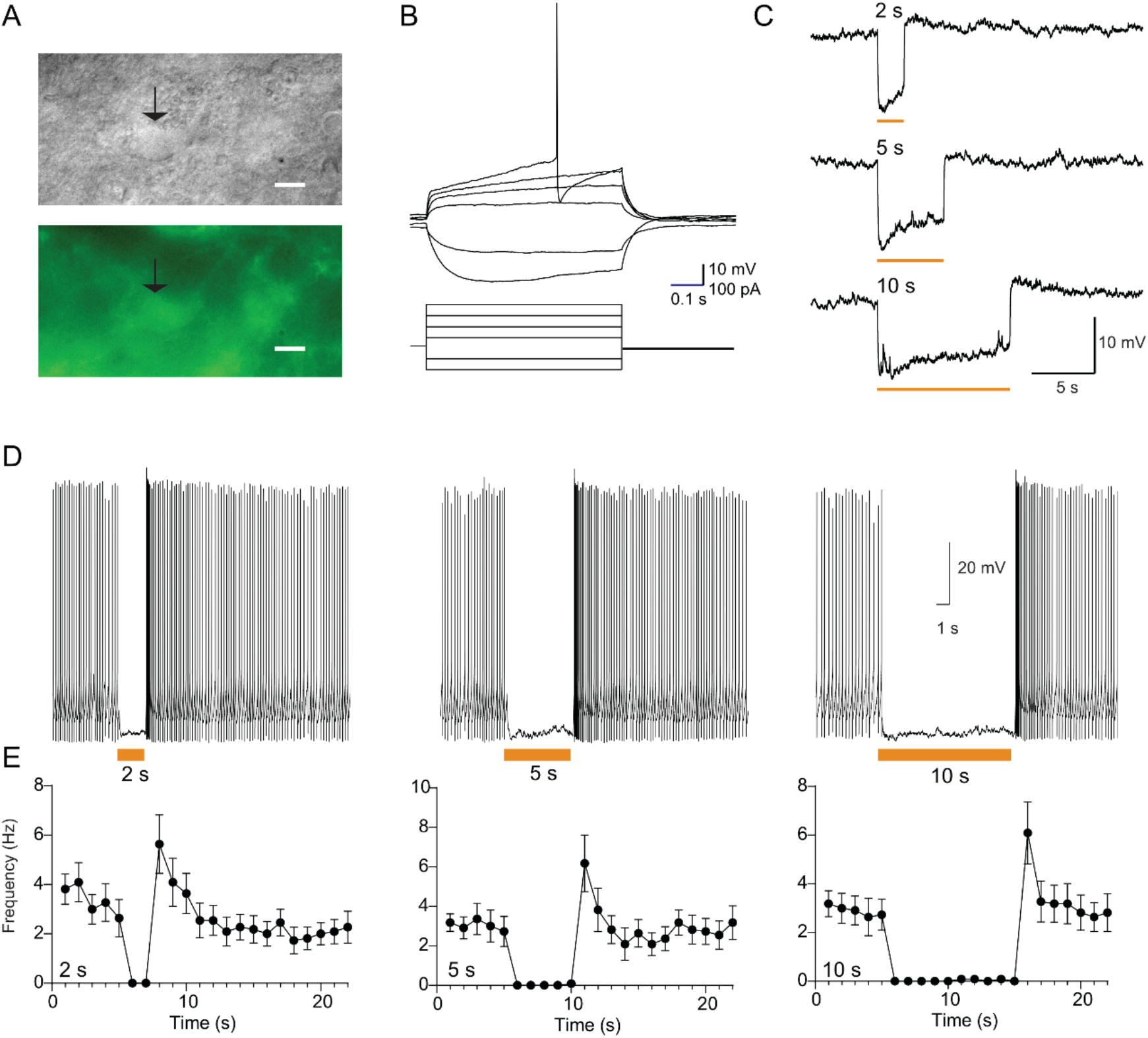
Whole-cell patch-clamp recordings from eNpHR3.0-positive VTA neurons. A, Gradient contrast and fluorescent image of an EYFP+ neuron in the VTA (indicated by arrow). Scale bar indicates 10 μm. B, Voltage response to hyperpolarizing and depolarizing current injections. C, Typical light-evoked hyperpolarisation, to 2, 5, and 10 s orange light pulses. D, Representative response of a spontaneously firing neuron before during and after 2, 5, and 10 s light pulses. E, Summary data plot of mean (± SEM; n = 11) firing frequency before, during, and after photoinhibition. Orange bars indicate timing of light presentation.

### Experiment 2: Contingency and duration sensitivity of photoinhibition

#### Histology

Figure 2A shows the location of viral infusion (each animal at 20% opacity) and cannulae tips in the VTA for TH::Cre+ and TH::Cre− animals. Five rats had misplaced or lost cannuale and were excluded from all analyses, leaving 10 TH::Cre- and 9 TH::Cre+ animals included in the final analyses.

**Figure 2.**
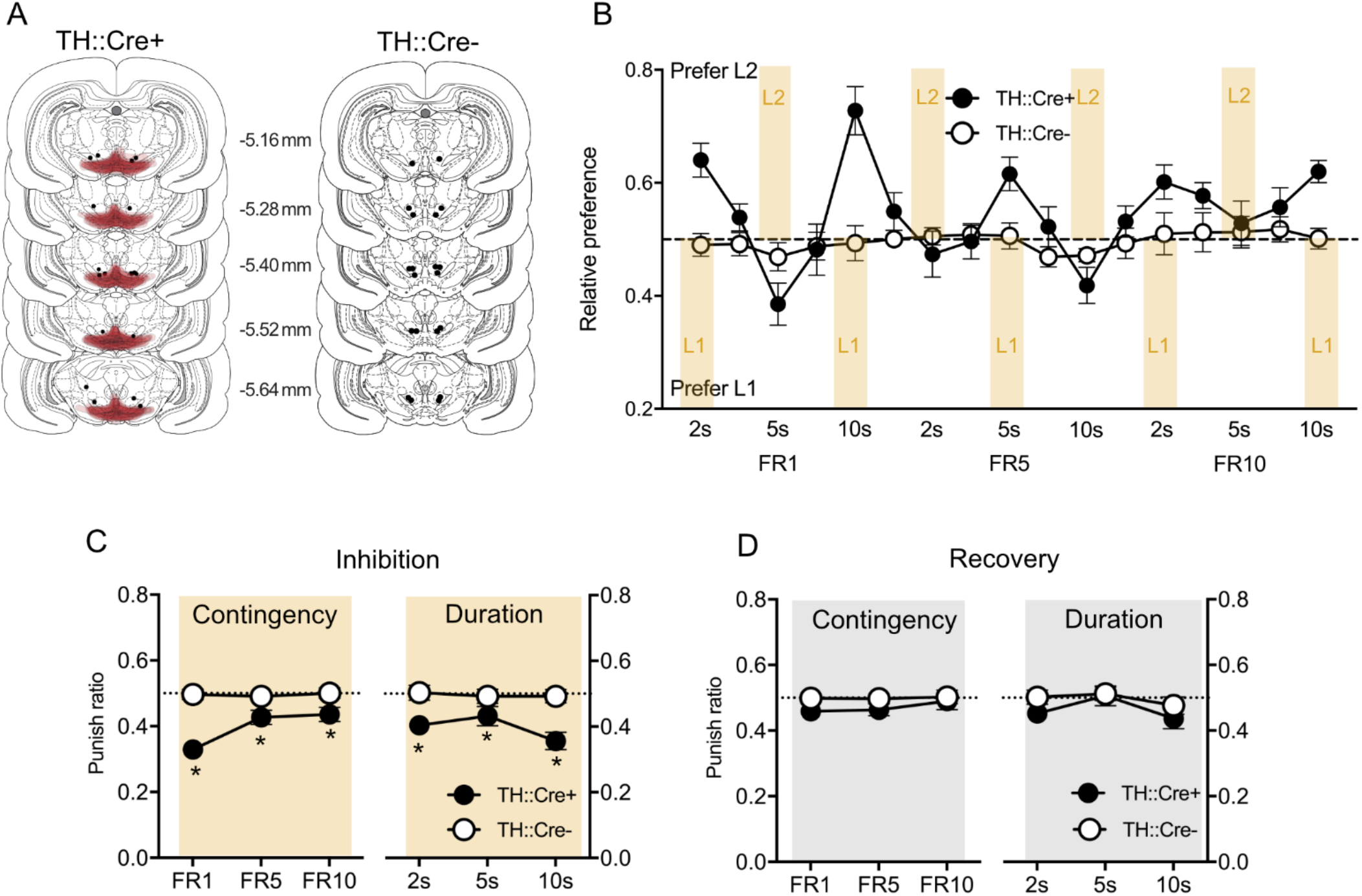
A, Location of AAV expression (each rat at 20% opacity) and fiber optic tips in the VTA. All distances in mm from Bregma. B, Mean and SEM preference for Lever 1 (L1) and Lever 2 (L2) across days in Experiment 2. Yellow shaded regions indicate photoinhibition days and identity of photoinhibited lever. C, Mean and SEM punishment suppression ratios across variations in instrumental contingency and photoinhibition duration. D, Mean and SEM punishment suppression ratios for recovery days across variations in instrumental contingency and photoinhibition duration. * p < .05.

#### Behavior

At the end of training, there were no overall difference between groups in lever presses (F(1,17) < 1; *p* > .05), no overall difference in responding on the to-be punished and to-be unpunished lever (F(1,17) < 1; *p* > .05) and no group × lever-interactions (F(1,17) < 1; *p* > .05).

Figure 2B shows lever preferences across the course of the experiment. These preferences are expressed as ratios whereby 0.5 indicates no lever preference, 1 indicates complete preference for lever 2 (L2) and 0 indicates complete preference for lever 1 (Lever 1). Photoinhibition was delivered following responses on one of the levers (indicated by yellow shading) followed by a recovery day with no photoinhibition. The identity of the photoinhibited lever alternated. So, these ratios track preference for the non-photoinhibited versus photoinhibited lever across photoinhibition schedules, durations, and across recovery. TH::Cre− rats showed no preference for either lever across days, but TH::Cre+ animals expressed consistent avoidance of the lever paired with photoinhibition, appropriately switching their preference away from the lever paired with photoinhibition each day regardless of the identity of that lever (L1 or L2).

The mean and SEM ratios during inhibition sessions are shown in Figure 2C. Overall, TH::Cre+ animals showed greater suppression of punished responding than TH::Cre− animals (F(1,17) = 30.82 *p* < .05). There was a main effect of schedule (F(1,17) = 12.16; *p* < .05), showing that the degree of suppression depended on the contingency of photoinhibition such that suppression significantly decreased as the contingency weakened. This interacted significantly with group (F(1,17) = 10.38; *p* < .05), confirming contingency sensitivity was related to photoinhibition, and follow-up comparisons showed that TH::Cre+ animals showed significantly greater suppression than TH::Cre− animals at each ratio (F(1,17) = 26.08, 9.42, and 16.71, p < .05). There was also a main effect of stimulus duration (F(1,17) = 6.49; *p* < .05), showing that longer photoinhibition was more effective than shorter photoinhibition at suppressing responding. This did not interact significantly with contingency (F(1,17) = 2.65; *p* > .05). Follow-up comparisons confirmed that TH::Cre+ animals showed significantly greater suppression than TH::Cre− animals at each photoinhibition duration (F(1,17) = 16.66, 11.34, and 28.43, *p* < .05).

The mean and SEM ratios during recovery sessions after photoinhibition are shown in Figure 2D. The question of interest here was whether the suppression of responding in the TH::Cre+ group observed during inhibition sessions would persist into recovery sessions without photoinhibition. Overall, ratios were higher during recovery sessions than photoinhibition sessions (F(1,17) = 26.60; *p* < .05) and this increase in responding during recovery sessions was greater for TH::Cre+ than TH::Cre− groups (F(1,17) = 21.90; *p* < .05). However, there was no main effect of schedule from the previous session (F(1,17) = 2.93; *p* > .05) nor an interaction with group (F(1,17) = 1.72; *p* > .05). There was also no significant main effect of photoinhibition duration from the previous session (F(1,17) = 4.05; *p* >.05) nor an interaction with group (F(1,17) < 1; *p* >.05).

### Experiment 3: Response learning versus stimulus learning

#### Histology

Figure 3A shows the location of viral infusion (each animal at 20% opacity) and cannulae tips in the VTA (only TH::Cre+ animals were used in this experiment). No rats were excluded from the analyses, so final group sizes remained Master (n = 8) and Yoked (n = 8).

**Figure 3.**
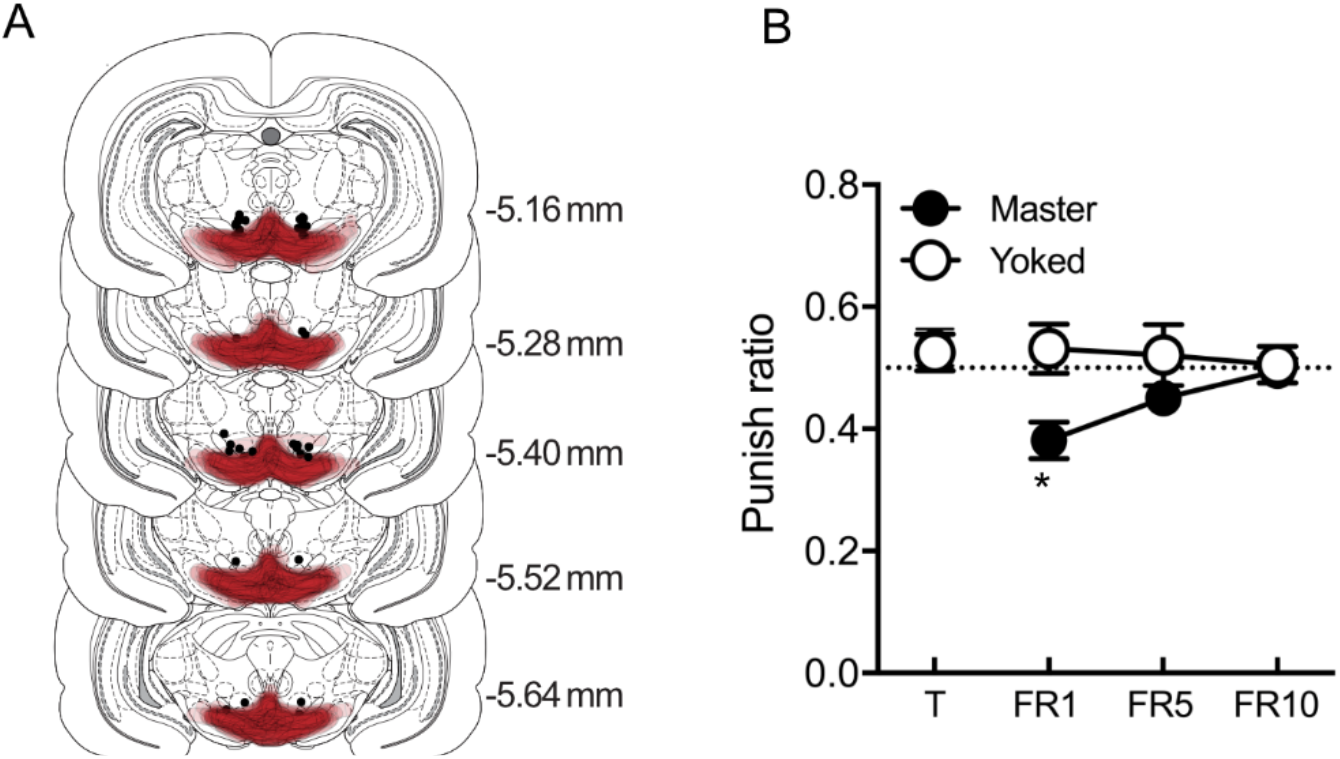
A, Location of AAV expression (each rat at 20% opacity) and fiber optic tips in the VTA. All distances in mm from Bregma. B, Mean and SEM punishment suppression ratios for Master and Yoked groups at the end of lever-press training (T) and across variations in instrumental contingency. * p < .05.

#### Behavior

At the end of training (T), there were no significant differences between Master and Yoked groups in lever presses (F(1,14) < 1; p > .05), no overall difference in responding on the to-be punished and to-be unpunished levers (F(1,14) = 1.39; p > .05) and no significant lever × group interactions (F(1,14) < 1; p > .05). The data of primary interest are the mean and SEM ratios from test, these are shown in Figure 3B. There was an overall significant difference between Master and Yoked groups (F(1,14) = 10.81; p < .05), showing significantly greater suppression in Master compared to Yoked groups. There was no overall significant effect of schedule (F(1,14) = 3.16; p > .05). However, there was a group × schedule interaction (F(1,14) = 8.10; p < .05). Follow-up analyses showed that Master rats were significantly suppressed relative to control rats at the FR1 (F(1,14) = 10.11; p < .05) but not FR5 (F(1,14) = 1.74; p > .05) or FR10 (F(1,14) = 0.06; p > .05) schedules.

### Experiment 4: Effects on instrumental extinction learning

#### Histology

Figure 4A shows the location of viral infusion (each animal at 20% opacity) and cannulae tips in the VTA for TH::Cre+ and TH::Cre− animals. Two rats had misplaced or lost cannuale and were excluded from all analyses, leaving 8 TH::Cre- and 6 TH::Cre+ animals included in the final analyses.

**Figure 4.**
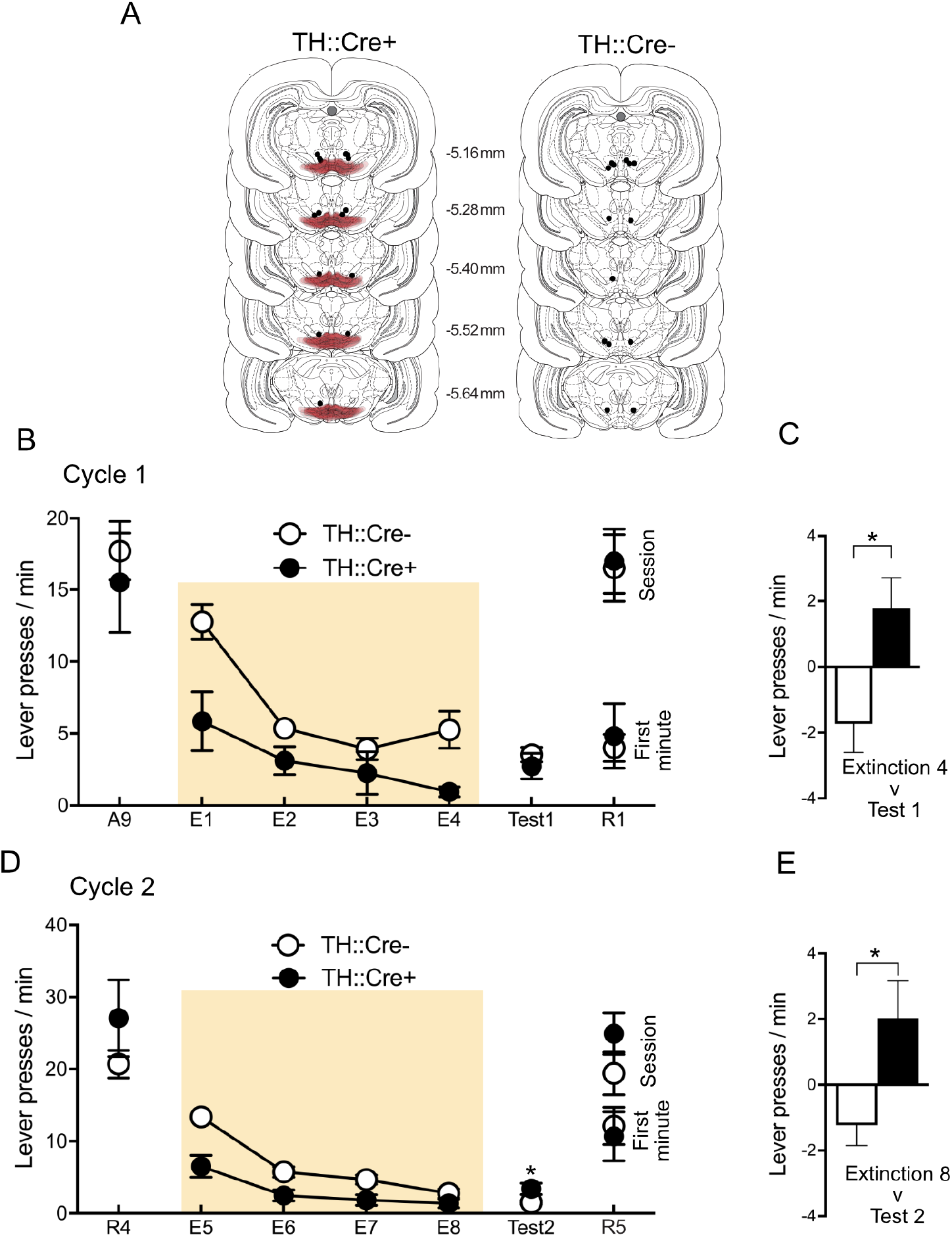
A, Location of AAV expression (each rat at 20% opacity) and fiber optic tips in the VTA. All distances in mm from Bregma. B, Mean and SEM lever press rates at the end of training (A9), across extinction training (E1 – E4), Extinction Test (Test 1) and Reacquisition (R1) during the first cycle of extinction training. C, Difference in lever press rates between Extinction 4 and Test 1 for TH::Cre+ and TH::Cre− rats. D, Mean and SEM lever press rates at the end of reacquisition training (R4), across extinction training (E5 – E8), Extinction Test (Test 2) and Reacquisition (R5) during the second cycle of extinction training. C, Difference in lever press rates between Extinction 8 and Test 2 for TH::Cre+ and TH::Cre− rats. * p < .05.

#### Behavior

There was no significant overall differences between TH::Cre- and TH::Cre+ groups in lever press rates at the end of training (A9) (F(1,12) < 1; p > .05). During the first cycle of extinction – test – reacquisition, there was a main effect of group across extinction (F(1,12) = 7.97; p < .05) showing that photoinhibition significantly reduced lever-pressing during extinction training. There was a main effect of day (F(1,12) = 39.91; p < .05), showing a significant decrease in responding across extinction training. However, there was no group × day interaction (F(1,12) = 1.79; p > .05). On test, in the absence of photoinhibition, there was no difference between groups (F(1,12) < 1; p > .05). However, whereas TH::Cre− showed a decrease in lever pressing between the last extinction session and test, TH::Cre+ animals showed an increase (interaction: F (1, 12) = 7.48, p < .05) (Figure 4C). There were no differences between groups in the initial reacquisition session (R1), either in the first minute (F (1, 12) < 1, p > .05) or across the complete session (F (1, 12) < 1, p > .05).

During the second cycle of extinction – test – reacquisition, there was again a main effect of group during re-extinction training (F(1,12) = 20.07; p < .05) showing that photoinhibition significantly reduced lever-pressing during re-extinction. There was again a main effect of day (F(1,12) = 50.05; p < .05), showing a significant decrease in responding across extinction training. There was also a group × day interaction (F(1,12) = 5.86; p < .05), showing that the decrease in responding across extinction was greater for TH::Cre− than TH::Cre+ group. On test, in the absence of photoinhibition, there was a significant difference between groups (F (1,12) = 6.58; p < .05), with TH::Cre+ animals showing significantly greater responding than TH::Cre− animals. Moreover, again, whereas TH::Cre− showed a decrease in lever pressing between the last extinction session and test, TH::Cre+ animals showed an increase (interaction: F (1, 12) = 7.39, p < .05) (Figure 4C). However, again, there were no differences between groups in the initial reacquisition session following this test (R5), either in the first minute (F (1, 12) < 1, p > .05) or across the complete session (F (1, 12) = 1.70, p > .05).

## Discussion

Here we asked whether inhibition of VTA dopamine neurons could serve as an instrumental punisher. Experiment 1 assessed whether response-elicited VTA inhibition caused specific suppression of that response, whether this suppression was affected by the contingency and duration of photoinhibition, and the degree to which effects persisted beyond inhibition sessions. The results showed that response-contingent photoinhibition of VTA dopamine neurons was sufficient to suppress instrumental lever-pressing, and that this was selective to the response earning photoinhibition because rats rapidly switched lever preferences across sessions to avoid photoinhibition. This selective suppression of responding is the same as that observed when footshock serves as the instrumental punisher (Bolles et al., 1980; Bolles et al., 1975). Suppression was also sensitive to both the contingency (F1 to FR10) and duration (2s to 10s) of photoinhibition. The stronger the Response–Photoinhibition contingency (e.g. FR1 v FR10) or the longer the duration of photoinhibition, the greater the response suppression. This profile of contingency sensitivity is the same as that observed when footshock is used as the instrumental punisher (Bolles et al., 1980). The response suppressive effects of photoinhibition were relatively transient, with no evidence that suppression persisted robustly beyond the photoinhibition sessions. Again, instrumental punishers such as mild footshock can also have transient effects on behavior (Church, 1963).

Nonetheless, these demonstrations are not themselves sufficient to identify instrumental response learning as the cause of response suppression. This is because the suppression observed in Experiment 2 might have been attributable to stimulus (i.e. Pavlovian) rather than response (i.e. instrumental) learning. For example, the rat was always in a specific location when it responded on the lever to earn photoinhibition and the presence of the lever itself was correlated with photoinhibition. These or other stimuli could have been learned as Pavlovian predictors of VTA dopamine neuron inhibition and so promoted avoidance of the lever and its location, causing a reduction in lever pressing. Indeed, photoinhibition of VTA dopamine neurons can support such place avoidance (Danjo et al., 2014; Ilango et al., 2014). Experiment 3 directly addressed this possibility using a yoking procedure. TH::Cre+ rats received photoinhibition of VTA dopamine neurons in the presence of one lever and but not the other. However, the groups differed in whether this photoinhibition was contingent on their lever-press behavior (Master group) or simply the presence of the lever they were pressing (Yoked group). Under these conditions, only animals trained under the instrumental contingency showed suppression of lever-pressing. As in Experiment 2, this response-dependent suppression was contingency-sensitive and selective to the photoinhibition-earning response. This profile of greater suppression relative to yoked controls is the same as that observed when footshock is used as the instrumental punisher (Boe & Church, 1968; Bouton & Schepers, 2015; Hoffman & Fleshler, 1965). Crucially, it suggests the suppressive effect of photoinhibition depended on the instrumental, not Pavlovian, contingencies in the task and that photoinhibition of VTA dopamine neurons is sufficient to support response learning.

Although Experiments 2 and 3 showed that response-dependent photoinhibition of VTA dopamine neurons selectively reduces responding on an instrumental basis, it does not readily explain how. It is unlikely that photoinhibition of VTA dopamine neurons achieved its effects by simply interfering with detection or processing of the reward, because the appetitive reinforcer and photoinhibition were delivered on different (interval versus ratio) schedules (Jean-Richard-Dit-Bressel et al., 2018) and photoinhibition was as effective in the presence of food reward as it was in the absence of this reward (Experiment 4). However, photoinhibition may have suppressed responding because it signaled an artificial reduction in or the absence of reward. Indeed, in Pavlovian preparations, brief optogenetic inhibition of VTA dopamine neurons at CS offset generates negative prediction error to mimic overexpectation learning (Chang et al., 2016) whereas excitation at the time of US omission prevents extinction learning (Steinberg et al., 2013). Response-contingent inhibition of these neurons could have reduced instrumental responding by mimicking signals of response non-reinforcement, i.e. acting as signals for instrumental extinction (Bouton et al., 2020). It follows from this possibility that photoinhibition of VTA dopamine neurons should serve to augment or deepen instrumental extinction learning, in the same way that augmentation of response or stimulus error during extinction training deepens extinction learning (Leung & Corbit, 2015; Rescorla, 2006).

Experiment 4 examined this. The results showed that response-dependent inhibition of VTA dopamine neurons during instrumental extinction training did cause a more rapid reduction in lever pressing than extinction training alone. However, this reduction was not achieved by augmenting extinction learning *per se* because lever-pressing recovered significantly more on test, in the absence of photoinhibition, compared to controls that received extinction only. Extinction was actually less effective when paired with photoinhibition of VTA dopamine neurons. Moreover, photoinhibition during extinction training had no effect on the rate at which animals could subsequently re-learn instrumental responding. This pattern of results, observed twice, is inconsistent with the possibility that photoinhibition of VTA dopamine neurons served as an instructive signal for instrumental extinction learning. To be sure, the overall magnitude of this recovery of responding among TH::Cre+ rats on test was modest, but the conditions of testing were not optimized for higher levels of responding because the food reinforcer had been absent from the testing conditions for 4 days prior. The important point is that only if TH:Cre+ animals had responded less on test or during retraining would we be able to conclude that extinction learning had been accelerated or deepened.

Moreover, the fact there was even a recovery of responding in TH::Cre+ rats at test is inconsistent with the possibility that photoinhibition deepened extinction learning. This pattern of results is precisely that observed when footshock is used as an instrumental punisher during extinction (Estes, 1944). As noted by Estes (1944), the enhanced reduction in responding during extinction followed by a return of responding on tests suggests that the punishment contingency (in this case VTA inhibition) did not promote the extinction process, but instead separately suppressed behavior, a suppression that was released during test. A congenial interpretation is that the suppressive effects of photoinhibition summated with the effects of non-reinforcement during extinction training, reducing overall rates of responding during extinction and thus reducing exposure to the extinction contingency. The amount of extinction learning produced by non-reinforcement depends on exposure to this contingency – the greater the responding during extinction training the greater the extinction learning (Rescorla, 2001). TH::Cre+ rats therefore learned less extinction as shown by more response recovery on test relative controls. Regardless, VTA dopamine neuron inhibition did not function as a learning signal for extinction.

However, it might still be argued that photoinhibition deepened extinction learning but that the conditions under which such learning was tested were more different for the animals that had received photoinhibition than those that had not. For example, the visual cueing properties of the 625nm light used for photoinhibition could come to form part of the extinction context and the absence of these visual cues on test would generate a context switch between extinction training and test in a manner similar to more global context changes (Bouton et al., 2020). This context switch could serve to renew the otherwise deepened extinction learning (Bouton & Bolles, 1979). The lack of effect of photoinhibition on reacquisition is not helpful in assessing this possibility because the rate at which animals reacquire instrumental responding after extinction can be insensitive to contextual manipulation (Willcocks & McNally, 2011). However, removal of discrete, response-contingent or non-contingent cues that were present only during extinction training does not renew or increase an otherwise extinguished response (McNally, 2014; Willcocks & McNally, 2014), suggesting this is an unlikely cause for the return of responding at test observed here.

In conclusion, we show that optogenetic inhibition of VTA dopamine neurons is sufficient to serve as an instrumental punisher. Optogenetic inhibition of VTA dopamine neurons causes a response-specific and contingency-sensitive reduction in instrumental responding that does not promote instrumental extinction. These effects of optogenetic inhibition of VTA dopamine neurons on instrumental responding are similar to the effects of instrumental punishment using mild aversive events.

## Acknowledgments

These experiments were supported by grants from the National Health and Medical Research Council (GNT116414) and Australian Research Council (DP19100482) to GPM.

